# Systematic Identification of Protein Phosphorylation-Mediated Interactions

**DOI:** 10.1101/2020.09.18.304121

**Authors:** Brendan M. Floyd, Kevin Drew, Edward M. Marcotte

## Abstract

Protein phosphorylation is a key regulatory mechanism involved in nearly every eukaryotic cellular process. Increasingly sensitive mass spectrometry approaches have identified hundreds of thousands of phosphorylation sites but the functions of a vast majority of these sites remain unknown, with fewer than 5% of sites currently assigned a function. To increase our understanding of functional protein phosphorylation we developed an approach for identifying the phosphorylation-dependence of protein assemblies in a systematic manner. A combination of non-specific protein phosphatase treatment, size-exclusion chromatography, and mass spectrometry allowed us to identify changes in protein interactions after the removal of phosphate modifications. With this approach we were able to identify 316 proteins involved in phosphorylation-sensitive interactions. We recovered known phosphorylation-dependent interactors such as the FACT complex and spliceosome, as well as identified novel interactions such as the tripeptidyl peptidase TPP2 and the supraspliceosome component ZRANB2. More generally, we find phosphorylation-dependent interactors to be strongly enriched for RNA-binding proteins, providing new insight into the role of phosphorylation in RNA binding. By searching directly for phosphorylated amino acid residues in mass spectrometry data, we identified the likely regulatory phosphosites on ZRANB2 and FACT complex subunit SSRP1. This study provides both a method and resource for obtaining a better understanding of the role of phosphorylation in native macromolecular assemblies.

## INTRODUCTION

Protein phosphorylation is a key cellular regulatory mechanism offering a reversible switch by which cells can activate and inactivate proteins^1^. Phosphorylation has been identified to be responsible for such processes as cellular replication^2^, apoptosis^3^, and transcription^4,5^ by actions such as modulating protein conformation^6^ and interactions^7,8^. With the increasing sensitivity of liquid-chromatography tandem mass spectrometry (LC-MS/MS) the ubiquity of protein phosphorylation is starting to be revealed and it is currently estimated that 75% of the human proteome may be phosphorylated^9^. Correspondingly, there have been many high-throughput phosphoproteomic studies resulting in hundreds of thousands of observed phosphorylation sites across many different cell types, tissues, and perturbed states^10,11^. This has resulted in a plethora of known phosphoproteins and phosphosites, but we lack insight into the function of most of these sites, or even if they are functional, leading to the suggestion that many observed phosphosites might represent biological noise^12–14^.

Recently, there have been several studies focused specifically on systematically identifying functions of protein phosphosites. Several of these studies have used computational strategies to make predictions of functional phosphosites. These approaches include use of 3D structure scanning to predict the effect of phosphates on interactions^8^, use of site conservation to predict a site’s functionality^13,15,16^, and more recently the development of a machine learning classifier that uses a variety of biochemical and evolutionary features to determine the probability of a site’s functionality in the form of a functional score^17^. These methods and their associated functional score predictions have provided great resources for the field and have given a sense of scale of the biological role of phosphorylation in the cell. Experimental approaches for high-throughput identification of protein phosphosite functions have been largely lacking. More recently, a promising approach has been to use a modified thermal proteome profiling (TPP) approach where the TPP protocol is conducted in conjunction with phosphoproteomics to identify functional phosphosites^18–21^. This method has proven useful to directly determine whether a phosphosite is likely to be functional or not. However, it fails to identify the specific nature of the function of the site, and this problem can be similarly indicated in nearly all of the current methods for identifying functional phosphorylation in a high-throughput manner.

To this end, we sought to develop an approach to specifically identify protein phosphorylation that regulates protein interactions. This method uses <u>dif</u>ferential <u>frac</u>tionation (DIFFRAC^22^) of chromatographically separated control and phosphatase-treated native (non-denatured) cell extracts, as measured by protein mass spectrometry on each chromatographic fraction, in order to detect changes in protein assemblies. Additionally, we developed and used orthogonal scoring methods for ranking proteins with the most notable phosphorylation-dependent changes in their separation behaviour. Using this “phospho-DIFFRAC” method, we gathered information on global functional phosphorylation and identified novel phosphorylation-mediated interactions in HEK-293T cells. Lastly, we were able to identify specific phosphosites in our data likely responsible for mediating several of the interactions. The data from this study provide a resource for better understanding the roles of phosphorylation in protein interactions.

## RESULTS & DISCUSSION

### Identifying functional protein phosphorylation using differential fractionation

The phospho-DIFFRAC method is grounded in previous work on co-fractionation mass spectrometry (CF-MS)^23,24^. CF-MS identifies protein complexes by separating proteins in their native assemblies along a biochemical gradient using non-denaturing separations and utilizes the tendency of proteins in a complex to co-elute, as detected by protein mass spectrometry on the resulting biochemical fractions, in order to identify protein interactions and complexes. CF-MS was more recently adapted in the form of DIFFRAC to identify RNA-binding protein (RBP) complexes by looking for changes in elution patterns between a control and RNase-treated sample when both samples are separated by size on a size-exclusion chromatography (SEC) column^22^. The DIFFRAC approach has been effective in different biological systems, including different cell types and embryonic tissues^25^, and we realized that the method could be applied to identify regulatory mechanisms of protein interactions. To this end, we applied DIFFRAC to identify phosphorylation-mediated protein interactions.

Phospho-DIFFRAC works by chromatographically separating, e.g. as by a non-denaturing SEC, two samples of cell lysate, one of which (the control) has protein phosphorylation preserved through the addition of a phosphatase inhibitor, and the other of which (the treatment) has accessible phosphate modifications removed by incubation with a non-specific protein phosphatase, Lambda Protein Phosphatase (**Fig 1a**)^26^. Following separation, proteins in each resulting fraction are identified and quantified using mass spectrometry^27^, resulting in an abundance profile for every protein that captures its elution across each chromatographic separation. Phosphorylation-dependent changes in protein abundance or interactions can then be identified by comparing each protein’s elution profiles from the control and phosphatase-treated samples. For example, a protein gaining a high molecular weight peak in the phosphatase-treated sample compared to the control suggests phosphorylation could be destabilizing an interaction involving that protein. Alternatively, a protein gaining a low molecular weight peak in the phosphatase-treated sample might suggest the presence of interactions stabilized by phosphorylation.

**Figure 1.**
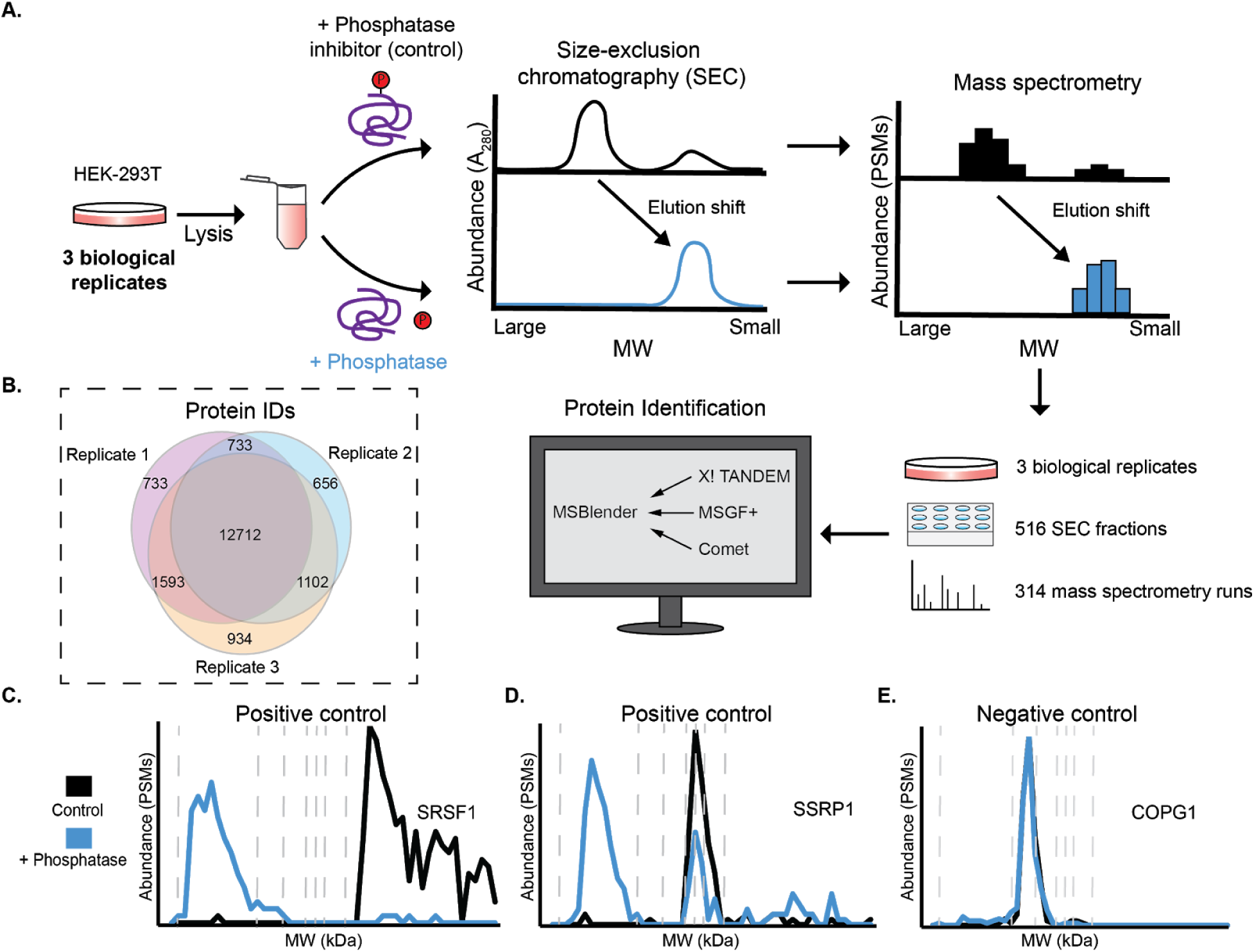
Workflow for phospho-DIFFRAC. A) Lysate from HEK-293T cells is separated into a control and phosphatase-treated sample. Proteins forming phosphorylation-dependent interactions will have elution shifts between the two samples. B) Venn diagram of protein observations for each biological replicate. C) Elution trace for serine/arginine-rich splicing factor 1 (positive control). Dashed vertical lines are molecular weight standards of 2000, 669, 443, 200, 150, 66, and 29 kDa. D) Elution trace for Structure-specific recognition protein 1 (positive control). E) Elution trace for coatomer subunit gamma-1 (negative control).

In order to evaluate the utility and power of this approach, we performed phospho-DIFFRAC on three biological replicates of HEK-293T cells grown in standard laboratory conditions (**Fig. 1a**). In total, across the three replicates, we collected and analyzed 314 SEC fractions by mass spectrometry. Over 12,000 proteins were identified consistently across all three replicates (**Fig 1b**), indicating high coverage of the proteome by replicate measurements, and we computed elution profiles for each of these consistently observed proteins across the phosphatase-treated and control sample separations.

To validate the method we first inspected the elution traces of proteins known to have interactions regulated by protein phosphorylation. Serine/arginine-rich splicing factor 1 (SRSF1) is a key component of the spliceosome as well as involved in transcription regulation, mRNA stability and transport, and translation^28^. SRSF protein kinase 1 (SRPK1) sequentially hyperphosphorylates the Arginine and Serine rich (RS) domain of SRSF1 resulting in a decreased affinity for RNA^29^. We observed a distinct shift in our data corresponding to the expected phosphorylation sensitivity of SRSF1. When dephosphorylated we observed a peak eluting in the megadalton (MDa) range likely corresponding to the RNA-bound SRSF1 and correspondingly we observed a low molecular weight peak for our putatively phosphorylated sample likely corresponding to the monomeric form of SRSF1 (**Fig 1c)**.

Additionally, the FACT complex subunit Structure-specific recognition protein 1 (SSRP1) is another protein known to form phosphorylation-sensitive interactions that we observed to have an elution shift (**Fig 1d**). This subunit is known to non-specifically bind to DNA when dephosphorylated and loses affinity to DNA upon phosphorylation by Casein Kinase 2 (CK2) ^30^. We observed a MDa peak in the phosphatase-treated sample that is likely DNA bound FACT complex and in both the phosphatase-treated and phosphatase-inhibited samples we observed a lower molecular weight peak between 66 and 200 kDa that closely corresponds to the expected unbound FACT complex mass of about 201 kDa.

Finally, we looked at a negative control protein that we did not expect to have an elution shift between the two samples. While it is rare to identify an abundant protein that is not phosphorylated, coatomer subunit gamma-1 (COPG1) provided a likely negative control as it is not known to be heavily phosphorylated and the sites which are phosphorylated on the protein are not highly predicted to be functional (Ochoa *et al*. max functional score: 0.327)^17^. As well, it is only known to form interactions with other members of the coatomer complex of which the assembly and composition is not known to be phosphorylation-sensitive. Tellingly, we did not observe any elution change between the two samples confirming that COPG1 is not likely to have interactions regulated by protein phosphorylation (**Fig 1e**). Phospho-DIFFRAC proved to be highly-replicable at discriminating between proteins known to have interactions regulated by protein phosphorylation and those not likely to be regulated by the modification.

### Scoring phosphorylation-dependent elution changes

Confident that the experimental component of the method was working, we then turned towards systematically scoring the interaction changes we identified. In the previous work identifying RNA-binding proteins a single “DIFFRAC score” was estimated using a normalized L1-norm (**equation 1 and 2**)^22^, treating each complete elution trace as a single vector. Due to the global nature of the DIFFRAC score, it tends to be somewhat less sensitive to proteins which have significant abundance changes within only a single fraction. (**Fig 2a**). To this end, we developed a second method for scoring each fraction. In short, this approach estimates a differential abundance for each fraction in the protein elution between the control and phosphatase-treated sample, normalizing the extent of differential abundances as a Z-score as in ref. ^31^, then combining the per-fraction Z-scores to generate a per-protein score (**equations 3-5**). **Fig. 2b** illustrates this scoring method by plotting the fold change for each fraction in relation to the protein abundance in each fraction. In accordance with expectation, fractions with large fold-changes tend to have more extreme Z-scores, in a manner that scales as a function of protein abundance.

**Figure 2.**
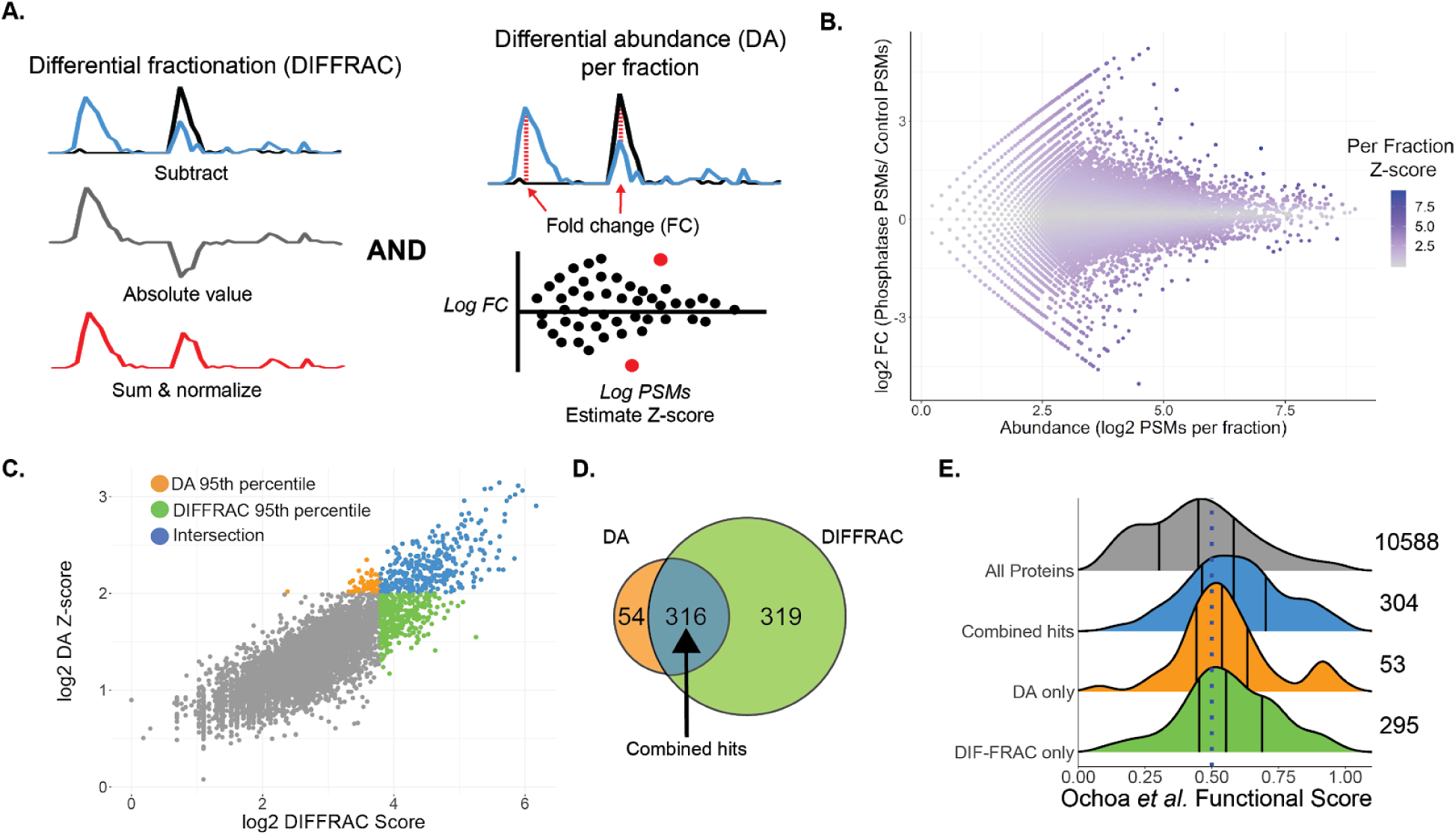
Scoring phosphorylation-dependent elution shifts. A) Two different approaches for scoring elution shifts. DIFFRAC scores changes by estimating an L1-norm for the entire elution trace as one vector, DA scores changes by estimating differences in each fraction between the two samples for each protein. B) Differential abundance (DA) Z-score per fraction compared to fold change for each fraction (mean of 3 replicates). C) Relationship between DIFFRAC and DA scores. The 95th percentiles of each scoring method are highlighted as well as the intersection of the two. D) Intersection between the 95th percentile of the DA and DIFFRAC scores. Same counts as in C. E) Protein functional score distribution for all proteins with functional scores (grey), phospho-DIFFRAC hits (blue), and for top-ranked proteins that were either only in the 95th percentile based on the DA score (orange) or the DIFFRAC score (green).

With this orthogonal scoring method in hand, we then compared it with the original DIFFRAC score, and observed strong concordance between the two methods in their ranking of hits (**Fig 2c**). We applied a 95 percentile threshold to obtain the top-ranked proteins from each scoring approach and we noted that the top proteins from each scoring method were largely in agreement with each other (**Fig 2c**). 316 proteins were in the 95th percentile for both scoring methods which, for simplicity, we will henceforward refer to as our phospho-DIFFRAC hits (**Fig 2d**), *i*.*e*. the set of proteins exhibiting the strongest phosphorylation-dependent changes to protein assembly status. To get a better understanding of how our method performed in identifying protein interactions regulated by functional protein phosphorylation, we utilized a previous study that used a classifier to predict the probability of a phosphosite being functional^17^. We collapsed the functional phosphosite scores for each protein by taking the max score and plotted the distribution of the scores for our combined hits as well as the non-intersecting top-ranked proteins from each scoring method (**Fig 2e**). In relation to the underlying distribution of functional scores for all proteins, our phospho-DIFFRAC hits are markedly enriched for proteins with high-scoring functionally phosphorylation sites.

### Global analysis of phosphorylation-dependent interactions

To better understand the role of functional protein phosphorylation in the cell we analyzed the frequencies of gene ontology (GO) annotations among the identified 316 phospho-DIFFRAC hits^32^. This revealed the striking enrichment of nucleic acid-, and in particular, RNA-binding proteins in our hits (**Fig 3a**). Phosphorylation is known to regulate transcription factor binding^4^ central RNA processes such as splicing^11^, but we were nonetheless surprised by RNA-binding dominating the GO annotations in our study, as opposed, for example, to metabolic enzymes, also known to be extensively regulated by phosphorylation^33^. To gain more insight into this phenomenon, we searched for enriched protein domains (Pfam) in our identified hits^34^. Consistent with the GO annotation analysis, the most enriched protein domains were RNA-recognition motifs (RRMs) (**Fig 3b**). The role of phosphorylation in regulating the RNA binding of RRMs has been identified and analyzed in the context of different proteins that each show a different role that phosphorylation plays in RNA-binding^29^. For example, in the case of SRSF1, hyperphosphorylation of the protein results in its loss of affinity for RNA. With this in mind, we looked into whether the number of phosphosites on a protein had any impact on it being involved in phosphorylation-mediated interactions. By looking at the total number of annotated phosphosites for each protein in our hits and comparing that to the number of phosphosites in all other proteins identified in our experiments we noticed a significant elevation in the number of total phosphosites for proteins in our hits (**Fig 3c**). Hyper-phosphorylation is known to affect spliceosome assembly, as well as seen in neurodegenerative diseases such as Tau in Alzheimer’s patients, but this could suggest a more ubiquitous role of hyper-phosphorylation in regulating protein interactions than previously expected^35,36^.

**Figure 3.**
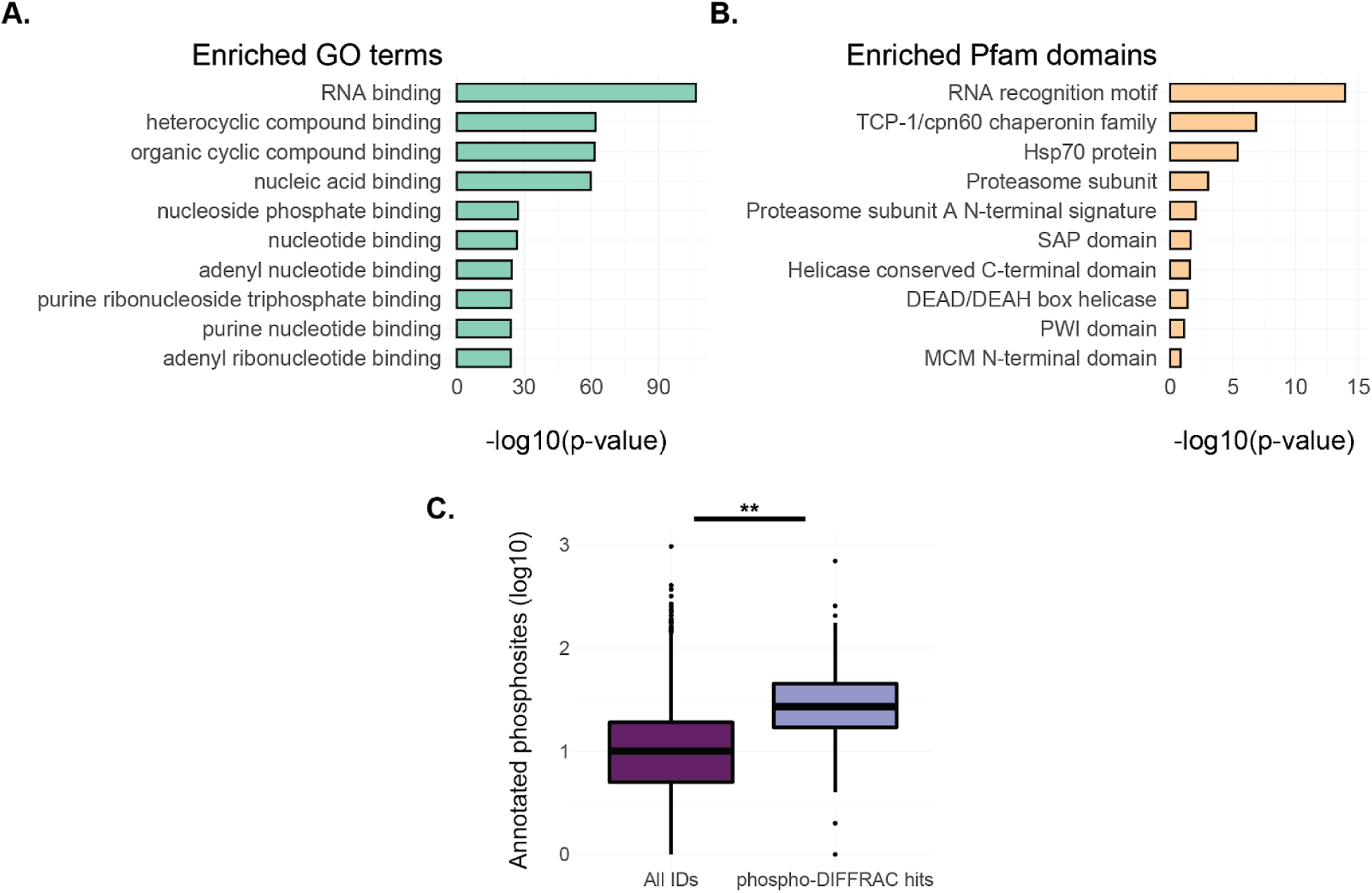
Global analysis of phosphorylation-dependent interactions. A) GO term enrichment for the phospho-DIFFRAC hits identified in this study. B) Enriched Pfam protein domains for the phospho-DIFFRAC hits. C) Boxplot of annotated phosphosite counts on proteins from PhosphoSitePlus (PSP) in all identified proteins (dark purple) and in phospho-DIFFRAC hits (light purple). Two-sided t-test t=-8.19, p value=6.44e-15.

### Phospho-DIFFRAC identified known and novel phosphorylation-dependent interactions

Using phospho-DIFFRAC, we were able to both recapitulate known phosphorylation-mediated interactions as well as identify novel ones. One novel interaction was identified in the homo-oligomeric protein complex of Tripeptidyl peptidase II (TPP2). TPP2 is a serine-protease associated with the proteasome that forms a homo-oligomeric megastructure with helices containing up to 36 subunits^37^. When comparing our phosphatase-treated sample to our control we consistently observed two distinct peaks in protein abundance; one of a high molecular weight between 669 and 2000 kDa consistent with the molecular weight of the homo-oligomeric form of the protein, and a smaller molecular weight peak eluting between 29 and 200 kDa consistent with the molecular weight of the monomeric form TPP2 around 138 kDa (**Fig 4a**). The peak corresponding to the monomeric molecular weight was either only observed in the phosphatase-treated sample or was observed in a higher abundance in the phosphatase-treated sample in each of the replicates. These data suggest a possible model in which dephosphorylation of the oligomerized complex results in disassembly of the complex down to the monomeric state (**Fig 4b**). To the best of our knowledge, this is the first evidence of a role for phosphorylation in the assembly of the megadalton TPP2 complex.

**Figure 4.**
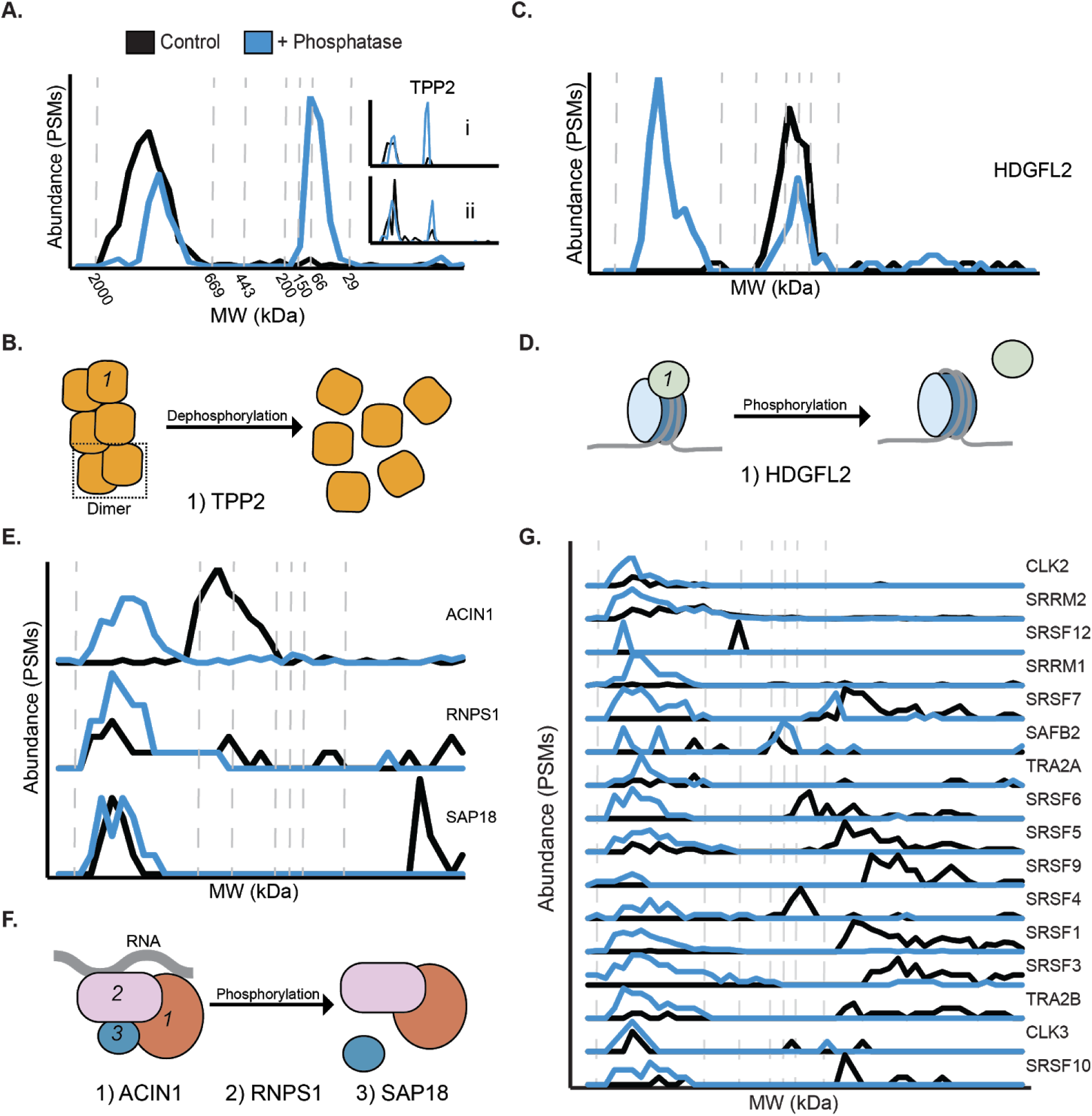
Phospho-DIFFRAC identified known and novel phosphorylation-dependent interactions. A) Elution trace of Tripeptidyl peptidase II. Black line is the control and the blue line is the phosphatase-treated sample. Inset i and ii are biological replicates. B) Model for TPP2 phosphorylation-dependent assembly. TPP2 dimer is boxed in the dotted line. C) Elution trace for Hepatoma derived growth factor-related protein 2 (HDGFL2). D) Model for HDGFL2 phosphorylation-dependent histone 3 nucleosome binding. E) Elution traces for subunits of the ASAP complex. From top to bottom: Apoptotic chromatin condensation inducer in the nucleus (ACIN1), RNA-binding protein with serine-rich domain 1 (RNPS1), Histone deacetylase complex subunit SAP18 (SAP18). F) Model for ASAP comlex phosphorylation-dependent composition. G) Elution traces for components of the spliceosome.

Another example of a novel phosphorylation-mediated interaction we identified is that of Hepatoma derived growth factor-related protein 2 (HDGFL2) and histone H3. HDGFL2 is a nucleosome binding protein that facilitates RNA polymerase II transcription by binding to H3K36me3 or, with less affinity, to H3K79me3^38^. HDGFL2 is a known phosphoprotein^39–41^, but there are no reports of phosphorylation affecting its interaction state. We identified two elution peaks for HDGFL2, one of a high molecular weight between 669 and 2000 kDa only observed in our phosphatase-treated sample and another at a lower molecular weight between 66 and 200 kDa that was observed in both samples (**Fig 4c**). This led us to the conclusion that the dephosphorylated form of HDGFL2 is capable of maintaining an interaction which the phosphorylated form is unable to keep. This interaction is likely with its known interaction partner H3C1. Our coverage of H3C1 was low, but the few spectral counts that were observed corresponded with the position of the higher molecular weight elution of HDGFL2 leading us to hypothesize that the interaction of the two proteins is phosphorylation-regulated (**Fig 4d, Supp Fig 1a**). The sensitivity of HDGFL2 to phosphatase-treatment indicates a functional role of phosphorylation in HDGFL2-regulated transcription and potentially in the cell proliferation observed in hepatocellular carcinoma samples^42^.

Additionally a regulatory role of phosphorylation in apoptosis was identified. The apoptosis- and splicing-associated protein (ASAP) complex is a RNA-binding complex, which is involved in mRNA processing and apoptosis regulation^43^. One of the subunits, RNPS1, is known to have it’s activity regulated by phosphorylation^44^, but is not reported to have interactions regulated by the modification. We observed RNPS1 and the rest of the ASAP complex to be phosphorylation-sensitive (**Fig 4e, Supp Fig 1b and 1c**). Interestingly, we observed a composition change in the complex between the phosphorylated and dephosphorylated samples. A slightly lower molecular weight complex between 200 and 669 kDa, consisting of at least ACIN1 and RNPS1 with the exclusion of the small SAP18 subunit was observed only in our control sample. The phosphatase-treated sample contained a higher molecular weight complex between 669 and 2000 kDa that contained all three complex members. (**Fig 4f**). While the full composition of the smaller complex is not currently known, it does indicate a role for phosphorylation in the assembly of this complex critical for cell apoptosis.

Finally, the spliceosome is known to assemble in a manner that is dependent on a complicated system of phosphorylation^45,46^. A high molecular weight form of the complex, between 669 and 2000 kDa, was only observed in the phosphatase-treated spliceosome (**Fig 4g**). This high molecular weight form closely corresponds to the intact spliceosome and indicates a role of phosphorylation in the maintenance of this complex. The spliceosome provides a clear example of the role that phosphorylation plays on macromolecules where the removal or addition can result in disassembly of the complexes. Additionally, it provides complex level support that the phospho-DIFFRAC method is working as expected.

### Identifying regulatory phosphosites in phospho-DIFFRAC data

We next sought to determine the specific phosphorylation sites regulating the interactions of some of our phospho-DIFFRAC hits. To do this we reanalyzed over 300 of our mass spectrometry experiments from this study using the proteomics software MSFragger to search for phosphorylated amino acid residues^47^. We then identified peptides in our phospho-DIFFRAC hits that were observed in both phosphorylated and dephosphorylated forms in the control and phosphatase-treated samples. Following this we were able to search for modified and unmodified peptides that showed differential elution (**Fig 5a**). Using this approach we identified phosphopeptides associated with elution peaks specific for either the control or the phosphatase-treated sample.

**Figure 5.**
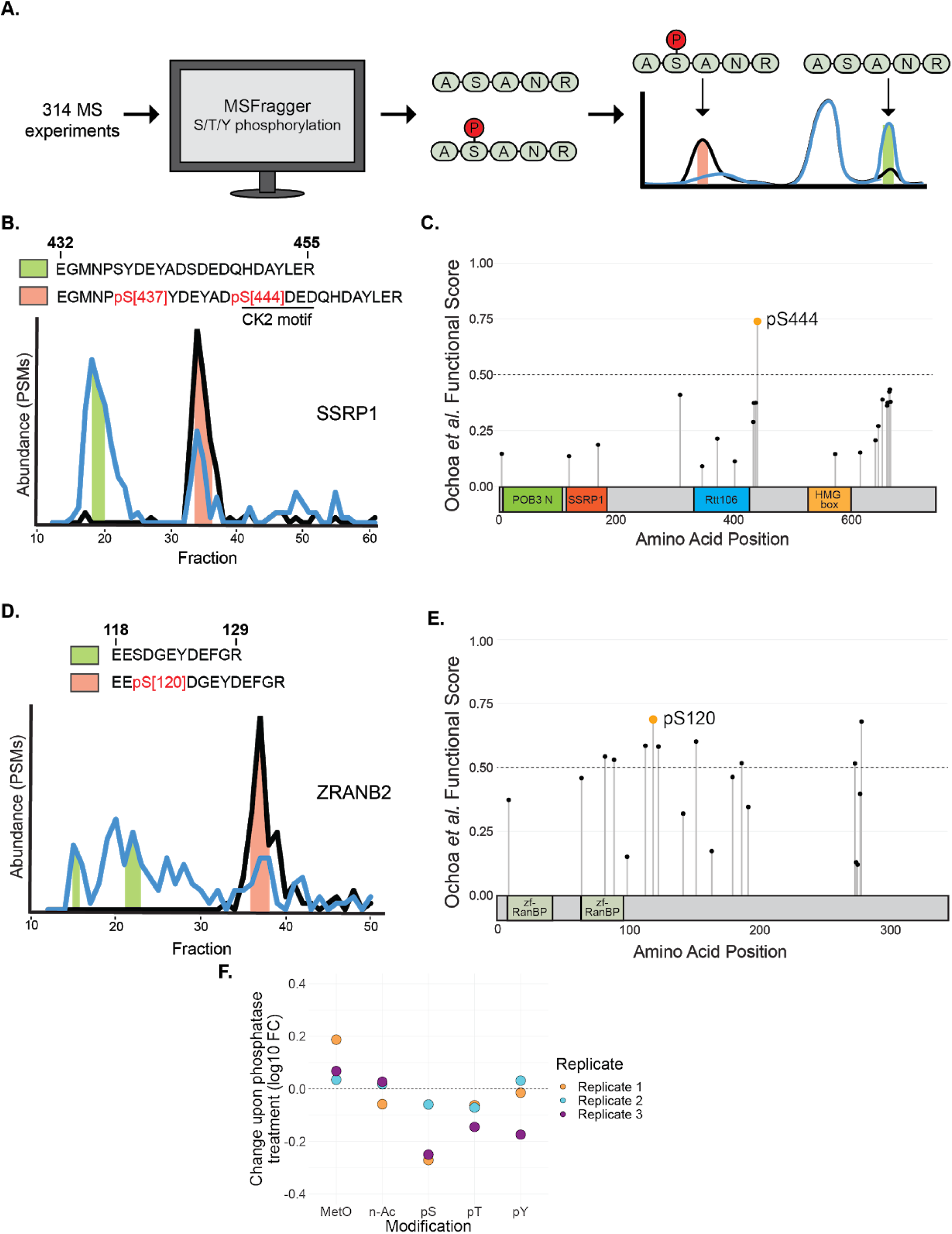
Differential phosphorylation analysis of phospho-DIFFRAC data. A) Workflow of differential phosphorylation analysis. Phosphopeptides were identified using MSFragger and peptides observed in both phosphorylated and dephosphorylated forms were analyzed to determine if the different modiforms eluted differently. B) Elution traces of SSRP1 with fractions that the differentially phosphorylated peptide was identified in shaded based on the phosphorylated form. Green shading indicates the dephosphorylated form and orange shading indicates the phosphorylated form. C) Functional scores for all phosphosites on SSRP1 overlayed on its domains. POB3 N: POB3-like N-terminal PH domain, SSRP1: Structure-specific recognition protein, Rtt106: Histone chaperone Rttp106-like, HMG box: High mobility group box. D) Elution traces of ZRANB2 with fractions that the differentially phosphorylated peptide was identified in shaded based on the phosphorylated form. Shading colors are the same as in B. E) Functional scores for all phosphosites on ZRANB2 overlaid on its domains. zf-RanBP: Zinc-finger in Ran binding protein. F) Difference in protein modification abundance between control and phosphatase-treated samples. MetO: Methionine sulfoxidation, n-Ac: N-terminal acetylation, pS: Serine phosphorylation, pT: Threonine phosphorylation, pY: Tyrosine phosphorylation.

One of these peptides came from Structure-Specific Recognition Protein 1 (SSRP1), a subunit of the heterodimeric FACT complex^48^. SSRP1 is known to non-specifically bind to DNA when dephosphorylated^30^. We identified two phosphorylated serine residues, at S437 and S444, that were associated with a distinct peak in the elution trace of SSRP1 (**Fig 5b**). To get a better idea of which site is more likely to be functional we examined the site-specific functional scores for all annotated phosphosites on SSRP1 (**Fig 5c**)^17^. Specifically, phosphorylated S444 is well observed in mass spectra in over 40 biological samples, is quantified in the top 5% of differentially regulated sites in 18 conditions, and is conserved throughout bilateria. Taken together, these data strongly suggest S444 is a functional phosphosite. Additionally, a previous study revealed that Casein Kinase 2 (CK2) is responsible for the phosphorylation that inhibits the DNA-binding ability of SSRP1^30^. As the site at S444 has a CK2 motif (SDXD), the combination of evidence suggests that this site plays a role in DNA-binding of the FACT complex^49^.

Additionally, we identified a potential regulatory site on Zinc finger Ran-Binding domain-containing protein 2 (ZRANB2). ZRANB2 is a component of the supraspliceosome containing a SR-like domain^50^. We identified a peptide on ZRANB2 that shows differential phosphorylation at S120 between the phosphatase-treated and control samples (**Fig 5d**). This phosphosite at S120 has not been previously indicated for regulating interactions with ZRANB2 and the spliceosome, but is observed as phosphorylated in mass spectra from over 50 biological samples and is conserved throughout Euteleostomi, all of which resulted in S120 having the highest functional score of all phosphosites on ZRANB2 (**Fig 5e**)^17^. This site thus seems responsible for the interaction between ZRANB2 and the spliceosome and serves as a foundation for future inquiry.

### Estimating the change in phosphorylation after phosphatase treatment

With the reanalyzed mass spectrometry data in hand, we were also able to make a larger survey of overall phosphorylation abundance changes between our two samples as compared to changes in other modifications (**Fig 5f**). Across all three replicates, we observed a decrease in Serine and Threonine phosphorylation, and a slightly less remarkable decrease in Tyrosine phosphorylation. This corresponds to the expected biases of Lambda Protein Phosphatase^51^. Interestingly, while N-terminal acetylation largely did not vary between the two samples, methionine sulfoxidation was observed to be more prevalent in our phosphatase-treated sample. The connection between phosphorylation and methionine sulfoxidation has been noted before with proteins that are phosphorylated tending to be proteins that undergo methionine sulfoxidation^52^.

### Comparison of experimental functional phosphorylation studies

Finally, we were able to evaluate the hits from our study in comparison to two previous thermal protein profiling (TPP) studies performed in human cells^19,20^. When looking at the functional score distributions, all three studies were observed to be enriched for functional phosphoproteins (**Supp Fig 2a**). Phospho-DIFFRAC was slightly lower in enrichment, likely because it identifies the interaction partners of functional phosphoproteins, which need not be functionally phosphorylated as well. Interestingly, the overlap in the significant hits identified in each study was reasonably large (e.g. as compared to interaction screens^53^), with the greatest intersection coming between phospho-DIFFRAC and the study of Huang *et al*.^19^ with nearly a third of the phospho-DIFFRAC hits being represented among the significant Huang proteins (**Supp Fig 2b**). We note that the extent of shared hits between this study and Huang is larger than that shared between the two TPP studies and remarkable considering the different methods used. Some of the disparity in shared phospho-DIFFRAC proteins with the two TPP studies might simply be a consequence of differing cell types (Huang *et al*.: HEK-293. Potel *et al*.: HeLa).

## CONCLUSIONS

In developing a new method for identifying phosphorylation-mediated interactions we have pushed forward the understanding of functional protein phosphorylation. The striking finding that many of the top proteins identified are RNA-binding proteins points to a more prominent role of phosphorylation in the assembly-dependent regulation of processes such as alternative splicing, transcription, and RNA transport than previously thought. Interestingly, our data suggest that phosphorylation is likely specifically regulating the interactions between proteins and RNA in addition to whatever other roles it may play in regulating protein activity. We expect that the data from this study will provide a source of future annotation of a new compendium of phosphosites. Finally, as the method is adaptable to assaying functional modifications in other cell types and species, it offers an opportunity to help in better defining cross-species functional phosphosites.

## Acknowledgements

The authors wish to thank Anna Mallam for discussion and constructive advice, Ophelia Papoulas and Dan Boutz for guidance with mass spectrometry, and Fan Tu for support with cell culture. This work was supported by grants from the NIH (R01 DK110520 and R35 GM122480 to E.M.M. and K99 HD092613 and LRP to K.D.) and the Welch Foundation (F-1515 to E.M.M.). Mass spectrometry data collection was supported by CPRIT grant RP110782 to Maria Person and by Army Research Office grant W911NF-12-1-0390.

## Data availability

All mass spectrometry data is available through PRIDE (Accession #PXD021422). Code used for the DIFFRAC and DA scores can be found at https://github.com/marcottelab/diffrac.

## Competing interests

The authors declare no competing financial interests.

## SUPPLEMENTARY INFORMATION

**Supplemental Figure 1.**
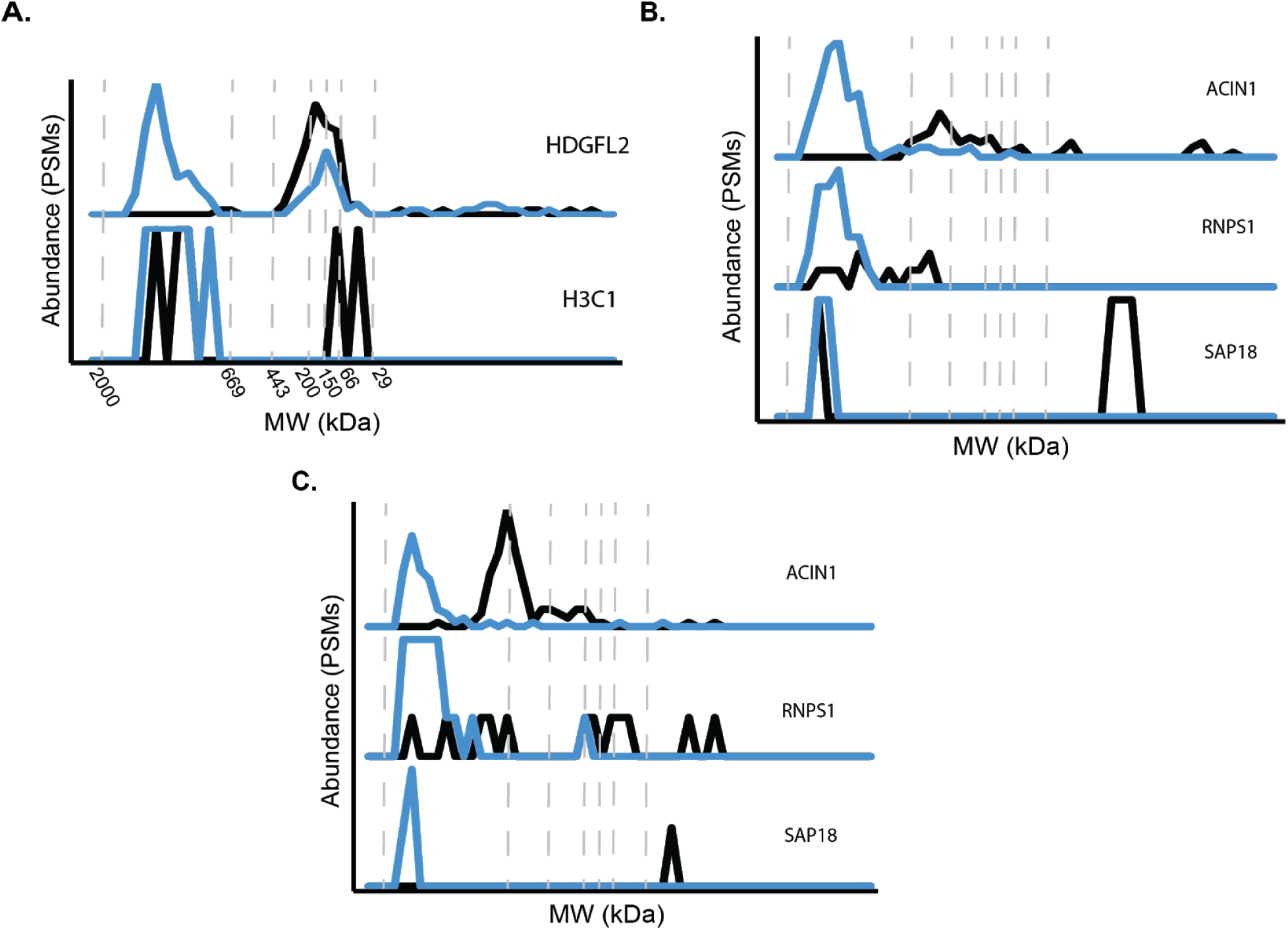
A) Elution traces for Hepatoma derived growth factor related protein 2 (HDGFL2), top, and histone 3 (H3C1), bottom. B and C) Elution traces for subunits of the ASAP complex from different biological replicates than that in Fig 4E. From top to bottom: Apoptotic chromatin condensation inducer in the nucleus (ACIN1), RNA-binding protein with serine-rich domain 1 (RNPS1), Histone deacetylase complex subunit SAP18 (SAP18).

**Supplemental Figure 2.**
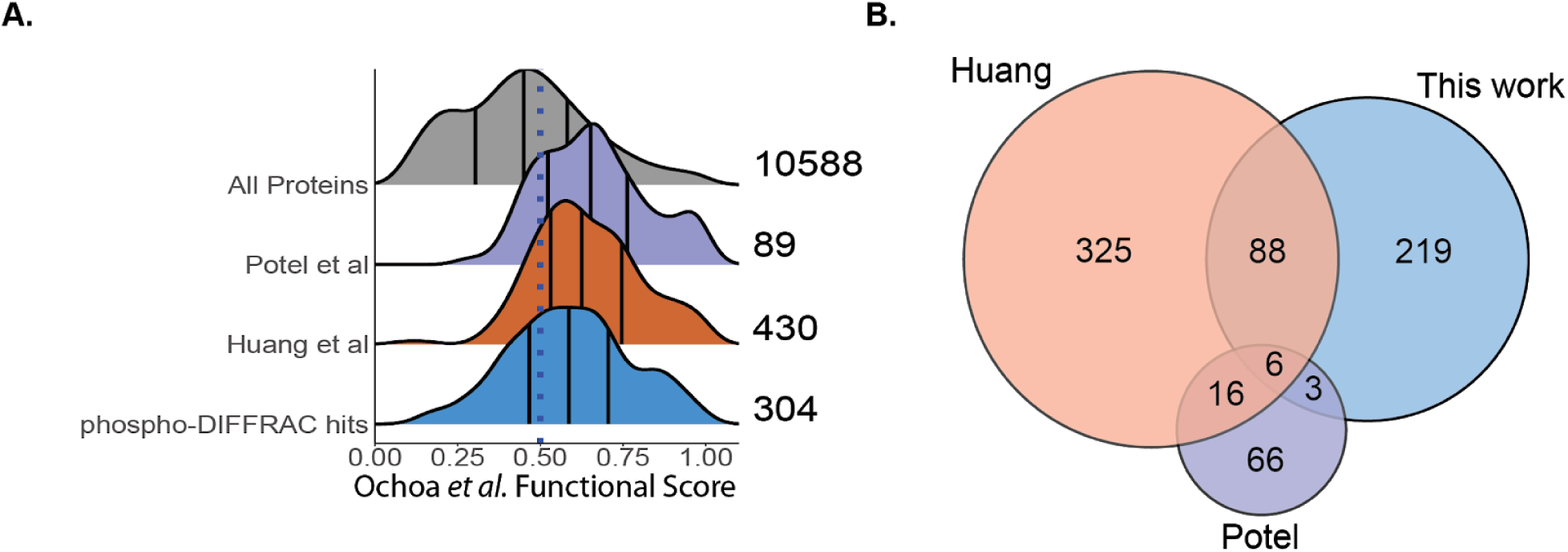
A) Protein functional score distribution for all proteins with functional scores (grey), hits from the study by Potel *et al*. (purple), hits from the study by Huang *et al*. (orange) and hits from this study (blue). B) Venn diagram of functional phosphoproteins identified in 3 different experimental studies.

**Supplemental Table 1**. Protein elution profiles for phosphatase-treated and control samples across all three replicates and fractions, reporting peptide spectral matches (PSMs) per protein per biochemical fraction.

**Supplemental Table 2**. DA z-score and DIFFRAC scores for phospho-DIFFRAC hits and all other proteins identified in the study, as well as per-fraction DA z-scores.

## MATERIALS & METHODS

### Cell culture

HEK-293T cells were grown in Dulbecco’s Modified Eagle Medium with 10% Fetal Bovine Serum at 37°C and 5% CO2. Cells were passaged between 70-80 % confluence.

### Phospho-DIFFRAC sample preparation

HEK-293T cells were washed and removed from 10 cm petri dishes using cold PBS pH 7.2. Cells were transferred to a 1.5 mL microcentrifuge tube and pelleted at 500 rpm for 3 minutes at 10°C. Supernatant was then removed and the wet cell pellet was weighed. 800 uL of Pierce IP lysis buffer (Thermo) was then added to the cells along with the addition of protease inhibitor (cOmplete ULTRA Tablets, Mini, EASYpack Protease Inhibitor Cocktail). The tube was kept on ice with soft intermittent mixing. The lysate was then spun at 17,000 g for 10 minutes at 4°C and the supernatant was split into two tubes. 100X phosphatase inhibitor (G-Biosciences PhosphataseArrest I) was diluted to 1X in one of the two tubes and PBS was added to the other. MnCl_2_ to a concentration of 1 mM and NEBuffer for Protein MetalloPhosphatases (NEB) diluted to a 1X concentration were added to both tubes. 4000 U of Lambda Protein Phosphatase was then added to the phosphatase tube and buffer was added to the other control sample. Samples were then incubated at 37°C for 30 min shaking at 60 rpm. Following incubation, samples were then filtered (0.45 μm Ultrafree-MC filter unit (Millipore)) spun at 12,000 g for 2 min, 4°C) to remove insoluble aggregates prior to fractionation.

### Size-exclusion chromatography

Treated and control cell lysates were subjected to size exclusion chromatography (SEC) using an Agilent 1100 HPLC system (Agilent Technologies, ON, Canada) with a multi-phase chromatography protocol as previously described^24^. Soluble protein (250 μL, 2 mg/mL) was applied to a BioSep-SEC-s4000 gel filtration column (Phenomenex) equilibrated in PBS, pH 7.2 at a flow rate of 0.5 mL min-1. Fractions were collected every 0.375 mL. The elution volume of molecular weight standards (thyroglobulin (Mr = 669 kDa); apoferritin (Mr = 443 kDa); albumin (Mr = 66 kDa); and carbonic anhydrase (Mr = 29 kDa); Sigma) was additionally measured to calibrate the column.

### Mass spectrometry

Fractions were filter concentrated to 50 μL, denatured and reduced in 50 % 2,2,2-trifluoroethanol (TFE) and 5 mM tris(2-carboxyethyl)phosphine (TCEP) at 55 °C for 45 minutes, and alkylated in the dark with iodoacetamide (55 mM, 30 min, RT). Samples were diluted to 5 % TFE in 50 mM Tris-HCl, pH 8.0, 2 mM CaCl2, and digested with trypsin (1:50; proteomics grade; 5 h; 37 °C). Digestion was quenched (1 % formic acid), and the sample volume reduced to ∼100 μL by speed vacuum centrifugation. The sample was washed on a HyperSep C18 SpinTip (Thermo Fisher), eluted, reduced to near dryness by speed vacuum centrifugation, and resuspended in 5% acetonitrile/ 0.1% formic acid for analysis by LC-MS/MS. Peptides were separated on a 75 μM x 25 cm Acclaim PepMap100 C-18 column (Thermo) using a 3-45 % acetonitrile gradient over 60 min and analyzed online by nanoelectrospray-ionization tandem mass spectrometry on an Orbitrap Fusion or Orbitrap Fusion Lumos Tribrid (Thermo Scientific). Data-dependent acquisition was activated, with parent ion (MS1) scans collected at high resolution (120,000). Ions with charge 1 were selected for higher-energy collisional dissociation fragmentation spectrum acquisition (MS2) in the ion trap, using a Top Speed acquisition time of 3-s. Dynamic exclusion was activated, with a 60-s exclusion time for ions selected more than once. MS from one of the three replicates was acquired in the UT Austin Proteomics Facility.

### Protein identification

Prior to protein identification, the human reference proteome (Acc: 02-2019) was downloaded from the Uniprot database^54^ to serve as the mass spectrometry reference database. Raw formatted mass spectrometry files were first converted to mzXML file format using MSConvert (http://proteowizard.sourceforge.net/tools.shtml) and then processed using MSGF+^55^, X! TANDEM^56^, and Comet^57^ peptide search engines with default settings. MSBlender^27^ was used to integrate peptide identifications and subsequently map to protein identifications. A false discovery rate of 1% was used for peptide identification. Protein elution profiles were assembled using unique peptide spectral matches for each protein across all fractions collected.

### Aligning elution traces

We observed by eye that across replicates there was a 1-2 fraction offset in elution traces. To mitigate against errors introduced by offsets in chromatography fraction positions, a set of proteins (VCP, FASN, GLU2B, COPG1, E41L2, WDR1, TERA, and PGK1) was selected as internal standards based on not exhibiting elution shifts upon addition of phosphatase and consistent observation in high abundance across all replicates. A sliding window Pearson coefficient was estimated to determine the appropriate integer fraction offset and the elution traces of all proteins were adjusted accordingly. Notably, the phosphatase-treated sample of replicate 3 had an offset of 12 fractions (due to collecting additional fractions during the flow-through) which was determined and adjusted in this manner. **Supp Table 1** reports the elution profiles with this offset applied.

### DIFFRAC score estimation

To score elution profile changes upon phosphatase treatment we compare a protein’s control elution trace to the phosphatase-treated elution trace, as illustrated in Figure 2A and first described in ref.^22^. To estimate this change we calculate the L1-norm between the two traces (**equation 1**),

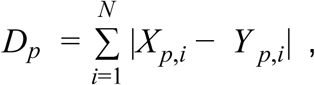

where N represents the total number of fractions collected and p represents an individual protein. X and Y represent abundance of control and experiment (phosphatase-treated) respectively. We next normalize *D*_*p*_ by the total abundance seen for protein *p* in both the control and treated group (**equation 2**),

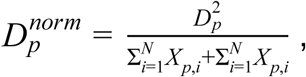

This was done for all three biological replicates and the median 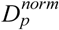 for each protein was used as our DIFFRAC score. This score can be converted to a Z-score or p-value to determine a significance threshold, as previously described^22^; for the purposes of this work, we simply considered proteins scoring above the 95th percentile as potential hits, *i*.*e*. those proteins with the strongest evidence for phosphosites regulating protein interactions. **Supp Table 2** reports all proteins identified with their scores.

### Differential abundance score estimation

To score elution profile changes based on peptide spectral matches per fraction (PSMs/fraction) we first estimated differential abundance between each control and phosphatase-treated fraction for every protein (**equation 3**),

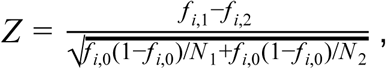

where *f* _*i*,1_ is the frequency of PSMs for a protein in the phosphatase-treated sample in fraction *i, f* _*i*,2_ is the frequency of PSMs for a protein in the control sample in fraction *i*, and the numerator represents the difference in sampled proportions of PSMs for protein in fraction *i* in the control and phosphatase-treated samples. The denominator represents the standard error of the difference under the null hypothesis in which the two sampled proportions are drawn from the same underlying distribution with the overall proportion *f* _*i*,0_ = (*n*_*i*,1_ + *n*_*i*,2_)/(*N* _1_ + *N* _2_) where *n*_*i*,1_ is the total PSMs for a protein in fraction *i* in the phosphatase-treated sample, *n*_*i*,2_ is the total PSMs for a protein in fraction *i* in the control sample, *N* _1_ is the total PSMs observed in the phosphatase-treated sample across all proteins and fractions, and *N* _2_ is the total PSMs observed in the control sample across all proteins and fractions. Z-scores were estimated for each fraction for all three replicates and combined to give a per-fraction Z-score, *Z*_*p*_, using Stouffer’s z-score method (**equation 4**):

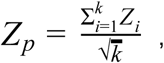

where *k* = 3. These scores were subsequently combined into a single protein Differential Abundance Z-score, *Z*_*DA*_, by applying Stouffer’s method to all per-fraction Z-scores for a given protein (**equation 5**):

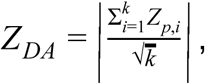

where *k* = # of fractions with observations.

### Gene Ontology and domain enrichment analysis

gProfiler was used for GO enrichment analysis. The 316 identified phospho-DIFFRAC hits were analyzed using the default gProfiler settings. The top 10 GO molecular function terms were used for visualization. Protein domain enrichment analysis was done using the DAVID tool on the phospho-DIFFRAC hits. Default settings were used for this analysis.

### Phosphopeptide identification

324 raw files from mass spectrometry experiments on both an Orbitrap Fusion and Lumos machines were converted to the mzML format using MSConvert with the peak picking filter selected. The files were split into their respective replicate directories and analyzed in 3 separate batches. MSFragger 2.4 on the command line was used for modification identification with the human proteome UP000005640 (accessed 3/25/2020) containing reversed protein sequences as a decoy database. A closed search was used with the fragment mass tolerance adjusted to 75 ppm. Variable modifications included in the search were N-terminal acetylation (42.01060), methionine sulfoxidation (15.99490), and serine/threonine/tyrosine phosphorylation (79.96633). Spectra were then assigned to peptides and processed using PeptideProphet with default settings. Protein identification was performed using ProteinProphet with default settings. A false discovery rate for all identification steps was set at 1%.

## Notes

### Competing Interest Statement

The authors have declared no competing interest.

